# Host sexual dimorphism affects the outcome of within-host pathogen competition

**DOI:** 10.1101/306985

**Authors:** Stephen A.Y. Gipson, Luis Jimenez, Matthew D. Hall

## Abstract

Natural infections often consist multiple pathogens of the same or different species. In multiple infections, pathogens compete for access to host resources and fitness is determined by how well a pathogen can reproduce compared to its competitors. Given the propensity for males and females to exhibit variation in pathogen-induced reduction in lifespan or fecundity, we explore how host sex may modulate the competitive ability of pathogens, potentially favouring the transmission of different pathogen genotypes. Using the *Daphnia magna - Pasteuria ramosa* model system, we exposed male and female hosts to either a single genotype infection or coinfections consisting of two pathogen genotypes of varying levels of virulence, measured as pathogen-induced reduction in host lifespan. We found that co-infections within females generally favoured the transmission of the more virulent pathogen genotype. Conversely, co-infections within male hosts resulted in equal transmission of competing genotypes, or favoured the transmission of the less virulent pathogen genotype in treatments where it established prior to the more virulent competitor. These results suggest that sex is a form of host heterogeneity which may influence the evolution of virulence within co-infection contexts and that one sex may be a reservoir for pathogen genetic diversity in nature.

## Introduction

Due to the ubiquity of pathogens in natural populations, individuals are usually infected with more than one type of pathogen or multiple strains of a single pathogen (Read and Taylor 2001; Rigaud et al. 2010; Balmer and Tanner 2011). In these multiple infection contexts, pathogens compete for access to host resources and the most fit competitor is often the one which induces the greatest impacts on host lifespan or fecundity (reviewed in Alizon et al. 2013). It is in this relationship between host exploitation and pathogen fitness that the predictions for multiple infections vary from those of single infections. While prudent pathogen exploitation leads to the highest pathogen fitness in single infection contexts, represented by a trade-off between pathogen reproduction and host harm (Bull 1994; Frank 1996; Alizon et al. 2009), a pathogen’s ability to acquire more resources than competing strains largely dictates relative pathogen fitness in multiple infections. Yet, host populations are rarely homogeneous and pathogens will encounter hosts that vary in characteristics such as age at exposure, nutritional background, or genotype (Wolinska and King 2009). Consequentially, the relative fitness (Hodgson et al. 2004; Izhar et al. 2015; Louhi et al. 2015) as well as the competitive exclusion or coexistence of co-infecting pathogens (de Roode et al. 2004) will depend not only on the virulence of the pathogens involved but also on the characteristics of their hosts.

A near ubiquitous source of host heterogeneity with the potential to modify the outcome of within-host pathogen competition are the differences between male and female hosts. Exhibiting differences in life-history strategies, physiology, or behaviour (Parker 2006; Schärer et al. 2012), the sexes vary in many characteristics that can influence pathogen exploitation (Duneau et al. 2012; Gipson and Hall 2016). For example, the sexes often vary in their relative immune investment (Rolff 2002; Zuk 2009) with one sex experiencing increased prevalence or severity of infection across many species (Poulin 1996; Schalk and Forbes 1997; McCurdy et al. 1998; Sheridan et al. 2000; Zuk 2009; Cousineau and Alizon 2014). The sexes may also represent hosts of varying exploitative potential due to differences in lifespan or physiological traits such as body size (Christe et al. 2007; Duneau and Ebert 2012; Thompson et al. 2017; Gipson and Hall 2018). Ultimately, these differences can result in infection having sex-specific effects on host fecundity or lifespan (see Table 1, Cousineau and Alizon 2014; Giefing-Kröll et al. 2015; Klein and Flanagan 2016), characteristics which drive within-host competition between pathogens (Alizon et al. 2013; Bashey 2015). Yet, it remains largely unexplored how differences between male and female hosts may influence the nature of within-host pathogen competition (but see Thompson et al. 2017) and potentially favour the transmission of different pathogen genotypes.

**Table 1:** Description of the four infection types used in this study. Single infections consisted of one genotype and co-infections consisted of genotypes C19 and C24 or C19 and C1.

| Infection type | Description | Dose at 5 days old |  | Dose at 12 days old |  |
| --- | --- | --- | --- | --- | --- |
|  |  | Geno. A | Geno. B | Geno. A | Geno. B |
| Simultaneous single infection | Individual was exposed once to a single pathogen genotype | 40,000 spores | - | - | - |
| Simultaneous co-infection | Individual was exposed once to two pathogen genotypes | 20,000 spores | 20,000 spores | - | - |
| Sequential single infection | Individual was exposed to a pathogen at day 5 and then to the same pathogen at day 12. | 20,000 spores | - | 20, 000 spores | - |
| Sequential co-infection | Individual was exposed to one pathogen at day 5 and then to a different pathogen at day 12. | 20,000 spores | - | - | 20,000 spores |

Within-host competition between pathogens can take a number of forms including: 1) *Exploitation*, where pathogens compete over a limited host resource pool; 2) *Apparent*, where the immune response stimulated by one pathogen indirectly inhibits competitors; or 3) *Interference*, where pathogen reproduction directly inhibits competitors either through prior establishment or competition over physical space (Mideo 2009; Balmer and Tanner 2011; Alizon et al. 2013; Bashey 2015; Cressler et al. 2016) - each form having different predicted outcomes for virulence evolution. Resource competition, for example, may cause pathogens to overexploit their host, leading to increased virulence (Chao et al. 2000; de Roode et al. 2005a); yet competing pathogens may interfere with one another or plastically adjust their rate of exploitation, leading to decreased virulence in these competition contexts (Massey et al. 2004; Choisy and de Roode 2010; Eswarappa et al. 2012). Inhibition may also extend to infections where one genotype establishes prior to the other; often favouring the earlier establishing pathogen regardless of their virulence characteristics (de Roode et al. 2005a; Eswarappa et al. 2012). Yet, as the most fit competitor will depend on the specific characteristics of its host (de Roode et al. 2004; Hodgson et al. 2004; Råberg et al. 2006; Ben-Ami et al. 2008; Ben-Ami and Routtu 2013; Izhar et al. 2015; Louhi et al. 2015), heterogeneity may be an important factor in determining the virulence of co-infections.

Within each form of pathogen competition exist unexplored opportunities for heterogeneity due to host sex to modulate the way pathogens exploit their host and influence the outcome of coinfection. The previously discussed effects of host sex on fecundity and lifespan, for example, may affect the relative competitive ability of pathogens by impacting on patterns of pathogen virulence. Additionally, physiological differences between the sexes may cause one sex to have less resources or physical space for a pathogen to exploit (Duneau and Ebert 2012; Thompson et al. 2017; Gipson and Hall 2018), potentially intensifying the competition between pathogens over host resources, or limiting the potential for pathogens to establish in previously infected hosts. The sexes may also vary in their immune response upon infection which may affect subsequent pathogen establishment. Given the propensity for males and females to vary in ways relevant to pathogen competition, sex may represent a further source of host heterogeneity impacting on pathogen transmission within co-infection contexts.

In this study, we use the *Daphnia magna - Pasteuria ramosa* model system to assess the impact that host sex has on the outcome of multiple infection. In this system, male *Daphnia* are smaller, exhibit shorter lifespans, are more resistant to infection, experience lower rates of infection-induced mortality, and allow for less production of pathogen spores; collectively suggesting that they are a more difficult resource to exploit than females (Duneau et al. 2012; Thompson et al. 2017; Gipson and Hall 2018). In nature, *Daphnia* may be infected with as many as eight genotypes of *P. ramosa* (Mouton et al. 2007) and have consequentially been used to study the evolution of virulence and competition between pathogens within co-infection contexts (Ben-Ami et al. 2008; Andras and Ebert 2013; Ben-Ami and Routtu 2013; Izhar et al. 2015). Co-infections in this system often resemble the virulence of the most virulent competitor in isolation and favour the transmission of more virulent genotypes except for situations in which the less virulent genotype establishes first (Ben-Ami et al. 2008; Ben-Ami and Routtu 2013). Yet co-infections play out differently between males and females, where females exhibit pathogen virulence and reproduction intermediate to that of the competing pathogens in isolation, whereas the outcome within males is more variable (Thompson et al. 2017). Remaining to be explored though is how these sex-specific patterns of co-infection may impact on the relative fitness among pathogen genotypes and the implications this may have for the maintenance of genetic variation and virulence evolution.

Using pathogen genotypes of known virulence characteristics (Clerc et al. 2015), we exposed genetically identical male and female *Daphnia* to either a single pathogen genotype or to a coinfection consisting of two pathogen genotypes of varying levels of virulence. We then varied the schedule of these exposures, either exposing hosts a single time, or allowing infection to establish prior to a subsequent exposure. We measured pathogen-induced reduction in lifespan (virulence), overall pathogen spore production (transmission), and the relative spore production of competing pathogen genotypes using microsatellite analysis. Due to the relationship between virulence and pathogen competitive ability, as well as the often observed sex-specific patterns of disease outcome, we make the following predictions: 1) the less exploitable sex will suppress variation in pathogen virulence, resulting in similar transmission of competing genotypes; 2) the more exploitable sex will allow for pathogens to exhibit variation in virulence, favouring the transmission of the less virulent genotype; and consequentially 3) the patterns of sequential coinfections in the more exploitable sex will resemble those of theory and empirical studies (de Roode et al. 2005a; Ben-Ami et al. 2008; Eswarappa et al. 2012), whereas the less exploitable sex will exhibit similar transmission of each genotype regardless of the relative virulence of the previously established genotype. We discuss the implications of these patterns for the maintenance of pathogen genetic variation and the evolution of virulence.

## Methods

*Daphnia magna* Straus is a globally distributed freshwater crustacean which produces genetically identical male and female offspring via cyclic parthenogenesis (Ebert 2005). During filter feeding, *Daphnia* encounter the bacterial pathogen *Pasteuria ramosa* which reduces the lifespan and fecundity of its host (reviewed in Ebert et al. 2016). *P. ramosa* is an obligate killing pathogen, transmitting exclusively horizontally after inducing host death. This experiment utilized host genotype HU-HO-2 and novel *P. ramosa* genotypes C19, C24, and C1. Prior to the experiment, we established a parental generation by isolating juvenile female *Daphnia* from pre-existing stock cultures and maintaining them in standardized conditions for three generations to minimize maternal effects. Juvenile female *Daphnia* were raised individually in 60-mL vials filled with 50 mL of artificial *Daphnia* medium (ADaM, Klüttgen et al. 1994; modified as per Ebert et al. 1998) and were transferred into fresh ADaM twice weekly. These females were maintained at 20°C, exposed to a 16-hour light to 8-hour dark cycle, and fed up to 5 million cells of *Scenedesmus sp.* green algae daily.

### Production of experimental animals

Once the third-generation standardized females released their first clutch, they were exposed to a short pulse of the hormone methyl farnesoate (300 μg/L, Product ID: S-0153, Echelon Biosciences, Salt Lake City, Utah) to stimulate the production of genetically identical male and female offspring. Following previously established methods (Thompson et al. 2017), the standardized females were transferred into 60-mL vials filled with 20 mL of hormone treated ADaM and were transferred into fresh hormone treated ADaM three times weekly. Male and female offspring were collected from the second and third clutches post hormone exposure. This treatment has previously been shown to have no detectible impact on host lifespan and fecundity, nor pathogen transmission and virulence (see Table 3, Thompson et al. 2017).

### Infection design

To measure the effect of host sex on the outcome of within-host pathogen competition, as well as how this competition proceeds when one genotype has already established, we randomly exposed males and females to either single infections or co-infections. These exposures were carried out in one of two exposure “schedules”: 1) *simultaneous* exposure occurred once when the individual was five days old or 2) *sequential* exposure occurred at 5 and 12 days old (see Table 1 for infection design details). All exposures consisted of a 40,000 pathogen spores. In sequential co-infections, the host was exposed to 20,000 spores of one pathogen genotype a week prior to 20,000 spores of the second to allow for prior establishment of infection. In sequential single infections, the host was simply exposed to 20,000 spores at each of the exposure periods by the same genotype. We herein refer to the multiplicity of genotypes and schedule of exposures collectively as “coinfection treatment.”

Three *P. ramosa* genotypes with previously studied disease characteristics were used in this study. When singly infecting female *Daphnia*, genotype C19 exhibits high virulence and low transmission (average infection duration: 45.29 days; average spore load: 8.56 million spores) whereas genotypes C24 and C1 cause similar infection outcomes, exhibiting lower average virulence and higher average transmission as compared to C19 (Clerc et al. 2015). Co-infections consisted of genotype C19 and either C24 or C1 as these genotype pairings represent similar virulence combinations and were thus predicted to exhibit similar patterns of competitive outcome. Additionally, genotypes C24 and C1 cannot be distinguished using our genetic analyses whereas C19 can be distinguished from C24 or C1, allowing us to determine the relative contribution of each pathogen genotype to the total spore production of co-infected *Daphnia* (see Genetic analysis section).

33 individuals of each sex were allocated to each co-infection or uninfected control treatment. In total this experiment consisted of 26 treatments (33 replicates * 2 sex * [3 simultaneous single infections + 3 sequential single infections + 2 simultaneous co-infections + 2 sequential coinfections treated with C19 first + 2 sequential co-infections treated with C19 second + 1 uninfected control treatment] = 858 individuals).

### Measures of disease characteristics

Survival was checked daily to assess lifespan of control and infected individuals. We calculated virulence by subtracting individual infected male or female lifespans from the average male or female control lifespan respectively. Upon host death, *Daphnia* were individually frozen in 500 μL of purified water for later determination of infection status and pathogen fitness as measured by overall production of transmission spores. Infection status was assessed by thawing a *Daphnia* sample, crushing it with a pestle, and noting the presence or absence of mature transmission spores using phase-contrast microscopy. Individuals identified as infected were immediately assessed for overall spore production using an Accuri C6 flow cytometer (BD Biosciences, San Jose, California). The spore load of each infected *Daphnia* was measured by diluting 10 μL of *Daphnia* sample into 190 μL of 5mM EDTA and loading this dilution into one well of a round-bottomed PPE 96-well plate. Custom gates based on fluorescence (FL3) were used to omit algae cells from the final count and custom gates based on side scatter (SSA) were used to identify only mature spores based on their distinct size and morphology compared to immature spores and animal debris (Ebert et al. 2016). Overall spore load was measured twice per individual and averaged.

### Genetic analysis and measures of within-host pathogen competition

To assess the fitness of co-infecting pathogen genotypes, we performed DNA extractions on coinfected *Daphnia* and determined the relative contribution of each pathogen genotype using variable number tandem repeats (Mouton et al. 2007). Pathogen genotypes were distinguished using primer sequences Pr1, Pr2, and Pr3 (Table 2, Mouton et al. 2007) which have been previously used to distinguish *P. ramosa* isolate P1 from isolates P4 and P5 (Ben-Ami and Routtu 2013) from which *P. ramosa* clones C19, or C24 and C1 are respectively derived (Luijckx 2012). As clones from isolates P4 and P5 cannot be distinguished using this method, co-infections always consisted of pathogen genotype C19 and C24 or C1.

**Table 2:** Summary of univariate and multivariate analyses of variance describing the effects of host sex, pathogen co-infection treatment, and their interaction on pathogen production of transmission spores, pathogen induced reduction in host lifespan and their multivariate interaction. Asterisks denote significant effects (α = 0.05).

| Multivariate ANOVA | Pillai's trace | Approx. *F* | d.f. | *p*-val |
| --- | --- | --- | --- | --- |
| **Production of transmission spores, reduction in host lifespan (days)** |  |  |  |  |
| Sex | 0.907 | 2939.443 | 2, 604 | <0.001 * |
| Pathogen co-infection treatment | 0.167 | 5.027 | 22, 1210 | <0.001 * |
| Sex : Pathogen | 0.108 | 3.133 | 22, 1210 | <0.001 * |
| Univariate ANOVA |  | *F* | d.f. | *p*-val |
| **Production of transmission spores** |  |  |  |  |
| Sex |  | 2048.260 | 1, 605 | <0.001 * |
| Pathogen co-infection treatment |  | 15.535 | 11, 605 | <0.001 * |
| Sex : Pathogen |  | 7.146 | 11, 605 | <0.001 * |
| **Reduction in host lifespan (days)** |  |  |  |  |
| Sex |  | 655.365 | 1, 605 | <0.001 * |
| Pathogen co-infection treatment |  | 6.110 | 11, 605 | <0.001 * |
| Sex : Pathogen |  | 0.526 | 11, 605 | 0.886 |

DNA extractions were performed using the EZNA Tissue DNA kit (Omega Bio-tek, Norcross, Georgia) with a modified protocol based on similar studies assessing the genetic composition of *P. ramosa* infections (Ben-Ami et al. 2008; Andras and Ebert 2013; Ben-Ami and Routtu 2013; Izhar et al. 2015). Immediately after spore counting, the crushed *Daphnia* samples were pelleted via centrifugation for 3 minutes at 12,205 RCF, supernatant removed, and washed with 1 mL of double-distilled water. The samples were again centrifuged using the same settings, supernatant removed, and suspended in 200 μL lysis buffer and 25 μL OB protease. The samples were then homogenized via bead beating with approximately 0.25 g of 0.1 mm zirconia beads for 2 minutes (1 × 10s, 1 × 20s, and 3 × 30s). Subsequently, samples were incubated in a heat block at 55°C for 1 hour before centrifugation at 10°C for 15 minutes at 5005 RCF. After collecting the supernatant, the DNA extraction proceeded as directed by the manufacturer protocol. An optional step of incubating samples for 2 minutes at 70 °C prior to elution greatly increased DNA yields. Final elution volume was 100 μL.

DNA was amplified via PCR with temperature cycling methods identical to Andras and Ebert (2013). Fragment analysis and genotyping was performed on these PCR products by AGRF (Melbourne, Australia) to determine the size of microsatellite alleles and the strength of their fluorescence (represented by peak height). The peak height ratio of the microsatellite markers was interpreted as the relative proportion of spores produced by each pathogen genotype as described by Ben-Ami et al. (2008); an approach that has also been used to quantify mixed sperm stores (Bussière et al. 2010). This proportion was multiplied by the absolute number of spores produced within the infected host to determine the relative transmission of competing genotypes.

### Statistical analyses

Only 823 of the 858 individuals initially set up for this experiment were used in final analyses as some individuals either died before infection status can be adequately determined (14 days post exposure, Clerc et al. 2015) or were removed due to experimental error. These 823 individuals consisted of 65 uninfected control individuals, 129 exposed but uninfected individuals, 319 individuals infected from single infection treatments, and 310 individuals infected from coinfection treatments of which *P. ramosa* DNA was extracted from 290 individuals. All statistical analyses were performed in R (version 3.4.1; R Development Team, available at www.r-project.org). Studies of multiple infection commonly explore how the most fit pathogen genotype is related to reduction in host fecundity or lifespan upon infection (reviewed in Alizon et al. 2013; Bashey 2015). Here, we focused on pathogen induced reduction in lifespan as a comparable measure of pathogen virulence between males and females. We then related patterns of virulence to overall pathogen transmission as well as the transmission of individual co-infecting pathogen genotypes within a mixed infection.

We first explored how the relationship between pathogen virulence and overall pathogen fitness changes due to host sex, pathogen co-infection treatment and their interaction using a multivariate analysis of variance (MANOVA Type III, *car* package, Fox and Weisberg 2011). Then we analysed the effects of sex, pathogen co-infection treatment, and their interaction on pathogen spore production and virulence using full-factorial analyses of variance (white corrected ANOVA Type III, *car* package). Using the *eemeans* package (https://github.com/rvlenth/emmeans) to perform post-hoc comparisons of the multivariate means for equivalence, we then explored how the relationship between virulence and overall spore production varied due to host sex or either of the two pathogen genotype combinations (C19 and C24 or C19 and C1). Finally, we tested whether the fitness of individual co-infecting pathogen genotypes changed due to host sex, co-infection treatment, or their interaction. To do this we fit a linear mixed model including each co-infection treatment (*lme4* package, Bates et al. 2015) with an individual’s unique ID fit as a random effect, an interaction between sex and pathogen genotype as fixed effects, and relative production of spores as the response. We also fit separate mixed models for each co-infection treatment in order to describe which co-infection treatment was driving patterns of significance in the full model.

## Results

### The effects of host sex on pathogen virulence, transmission, and their interaction

Our results indicate interactive effects of host sex and patterns of co-infection on the combined influence of pathogen fitness (the total spore production of co-infections or single genotype infections) and pathogen induced reduction in host lifespan (virulence). Across all single and coinfection treatments, the multivariate ANOVA reveals how infections in females lead to higher spore loads and a greater reduction in lifespan than in males (Fig. 1); but that the magnitude of any response to infection is not shared equally between the sexes due to the presence of the interaction term (Table 2). Figure 1 and Table 2 (univariate ANOVA) suggest that this interaction is largely driven by greater variation in spore loads between co-infection treatments occurring in females (Fig. 1a), with pathogens producing between 9.6 million and 5.9 million average spores within females and between 1.9 million and 1.1 million average spores within males. In contrast, females experienced higher levels of virulence than males, but the relative differences in virulence among co-infection treatments was similar between the sexes (Fig. 1b). Collectively this suggests that while pathogens which delay host death produce the greatest number of spores in females, this trade-off is more dampened in males where variation in virulence relates to a much smaller range in pathogen reproduction (Fig. 1).

**Figure 1:**
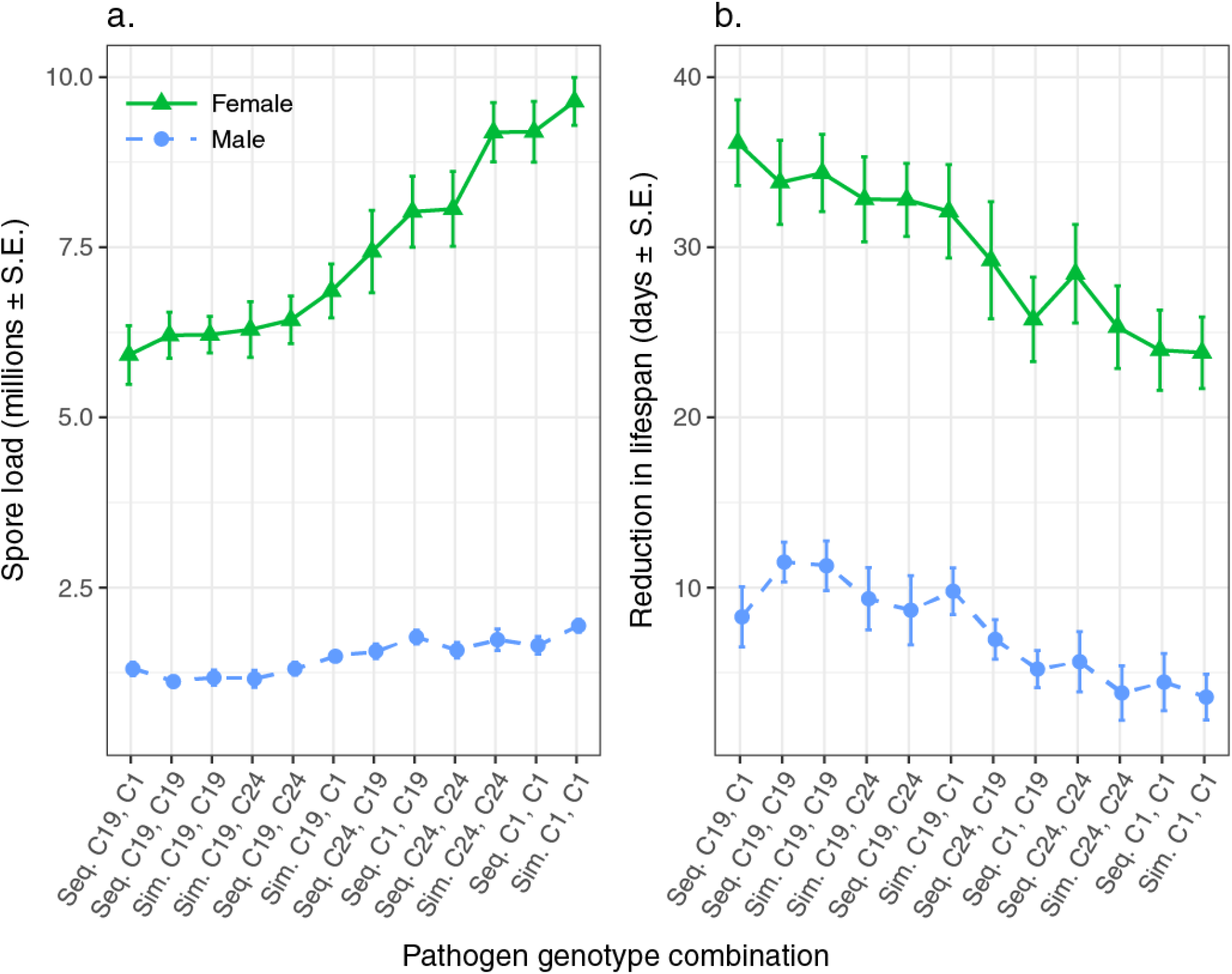
The influence of host sex and co-infection treatment on overall pathogen spore production (a) and reduction in host lifespan (b). Shown are treatment means and standard errors. Female results are represented by a solid green line with triangles and male results are represented by a dashed blue line with circles. Pathogen co-infection treatments are ordered by ascending average spore production in females. Sequential (Seq.) infection labels refer to the order in which the two pathogen genotypes established in their hosts whereas simultaneous (Sim.) infection labels refer to the two pathogen genotypes which established in their host at the same time.

Despite the difference in the scale of the trade-off between males and females, within each sex the relationship between pathogen transmission and virulence across co-infection treatments remained qualitatively similar. As expected, in females we found pathogen genotype C19 was more virulent and produced less transmission spores than C24 (Fig. 2a) or C1 (Fig. 2b) in single infection contexts. Similarly, sequential single genotype infections were indistinguishable from simultaneous single infections. Simultaneous co-infections always behaved like the more virulent pathogen in single infection contexts (C19), yet the order of sequential co-infections influenced the relationship between virulence and transmission differently based on the genotype of the competitor. In competitions with between C19 and C1, sequential co-infections always behaved like the genotype that established first (*i.e.* sequential C19, C1 behaved like simultaneous C19, C19; Fig. 2b). In contrast, the relationship between virulence and transmission of sequential coinfections were indistinguishable in competitions between C19 and C24 (Fig. 2a). Males exhibited comparatively similar patterns to females, with C19 more virulent and producing less spores than C24 (Fig. 3a) or C1 (Fig. 3b) in single infections; sequential single infections exhibiting similar patterns to simultaneous single infections, sequential co-infections with C24 exhibiting similar patterns regardless of infection order, and sequential co-infections with C1 behaving like the genotype which established first.

**Figure 2:**
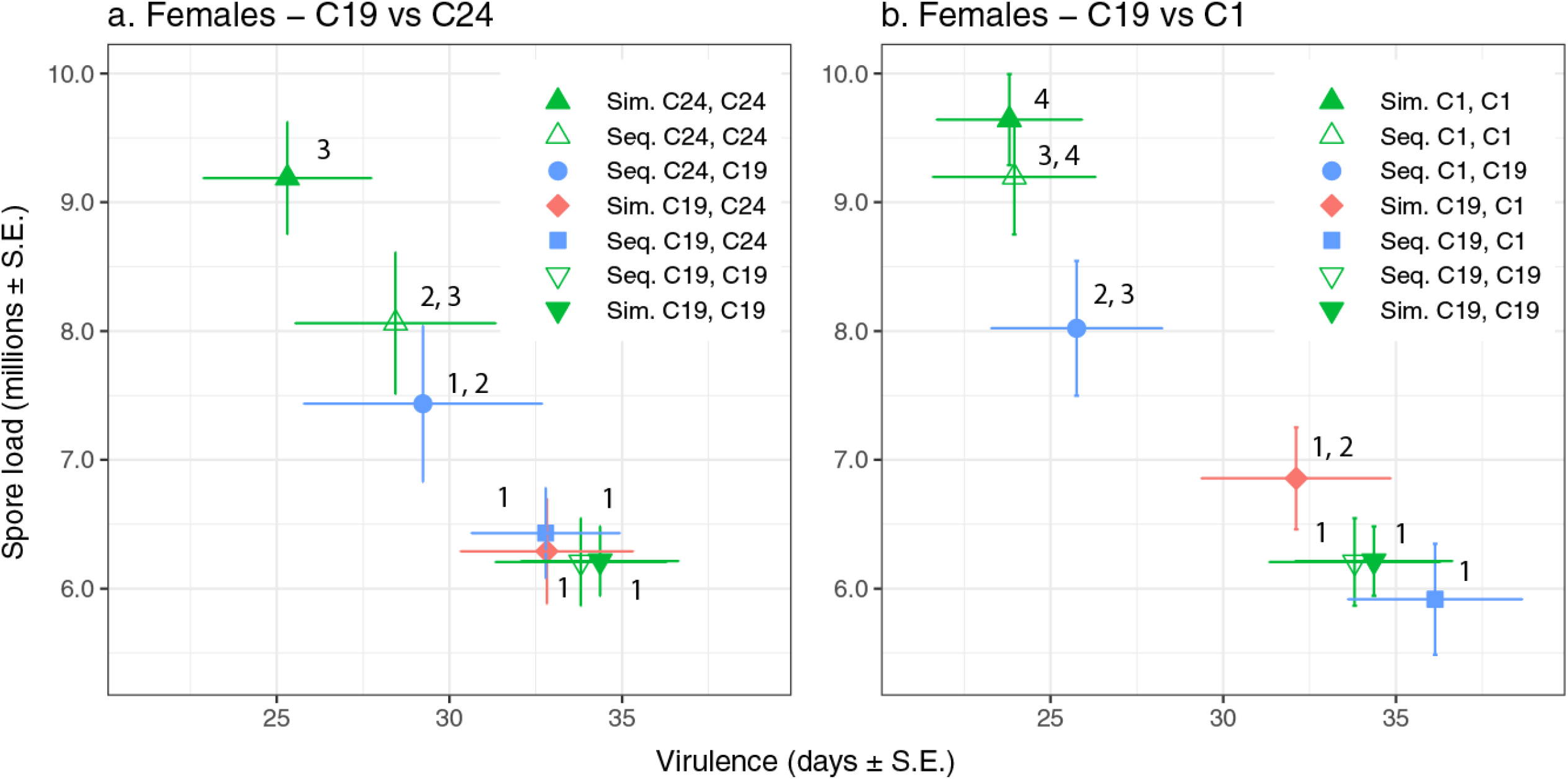
The influence of co-infection treatment on the relationship between overall pathogen spore production and reduction in host lifespan (virulence) for females exposed to pathogen genotype C19 and/or C24 (a) and females exposed to pathogen genotype C19 and/or C1 (b). Solid green triangles indicate single simultaneous infections, open green triangles indicate single sequential infections, blue squares or circles refer to sequential co-infections where the host was exposed to pathogen genotype C19 first or second respectively, and red diamonds refer to simultaneous coinfections. Shown are multivariate treatment means and standard errors with different associated numbers signifying a significant difference in multivariate mean between treatments. Sequential (Seq.) infection labels refer to the order in which the two pathogen genotypes established in their hosts whereas simultaneous (Sim.) infection labels refer to the two pathogen genotypes which established in their host at the same time.

**Figure 3:**
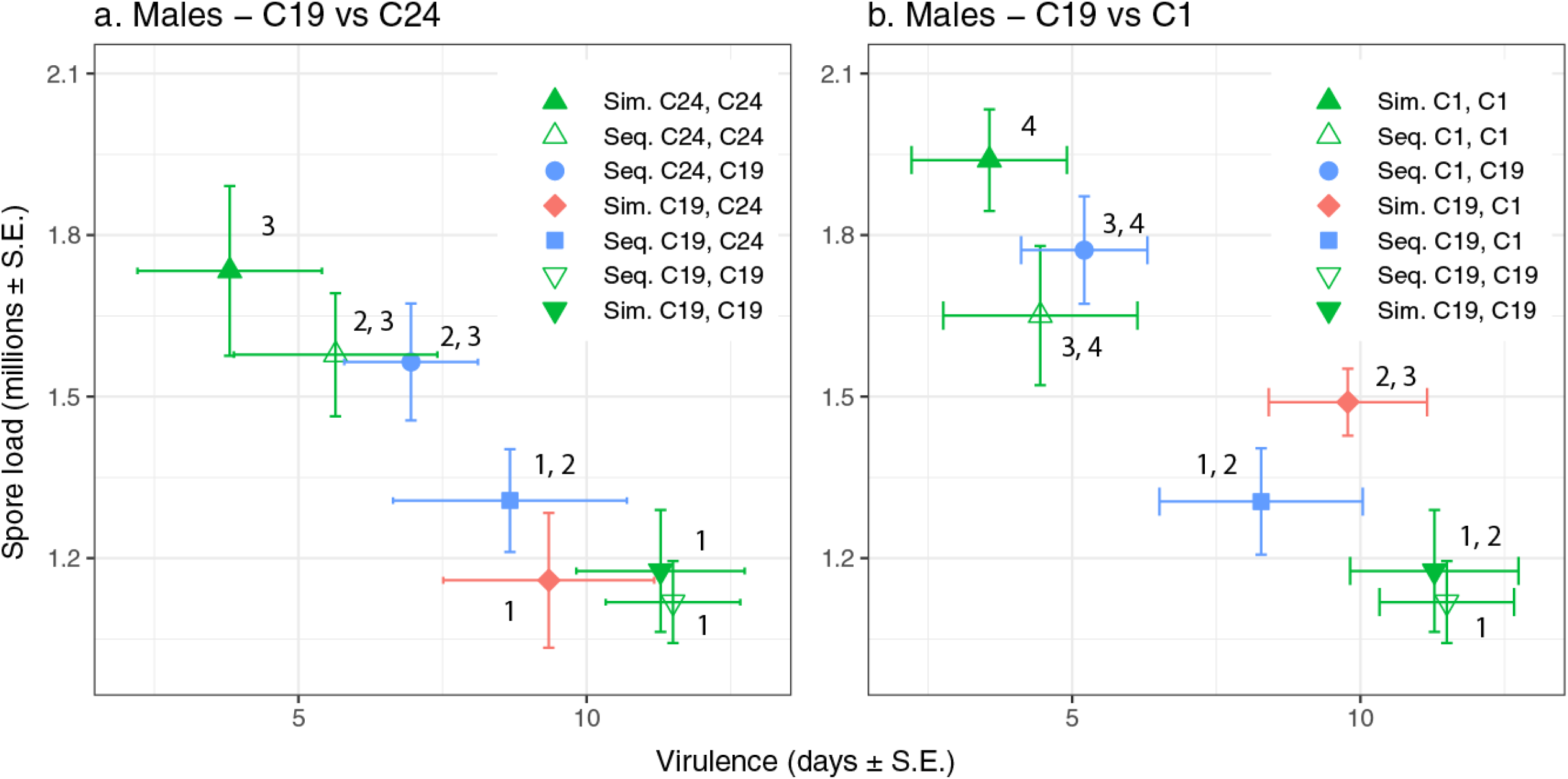
The influence of co-infection treatment on the relationship between overall pathogen spore production and reduction in host lifespan (virulence) for males exposed to pathogen genotype C19 and/or C24 (a) and males exposed to pathogen genotype C19 and/or C1 (b). Solid green triangles indicate single simultaneous infections, open green triangles indicate single sequential infections, blue squares or circles refer to sequential co-infections where the host was exposed to pathogen genotype C19 first or second respectively, and red diamonds refer to simultaneous coinfections. Shown are multivariate treatment means and standard errors with different associated numbers signifying a significant difference in multivariate mean between treatments. Sequential (Seq.) infection labels refer to the order in which the two pathogen genotypes established in their hosts whereas simultaneous (Sim.) infection labels refer to the two pathogen genotypes which established in their host at the same time.

### The effects of host sex on the relative fitness among co-infecting pathogen genotypes

Finally, we explored how host sex and pathogen genotype collectively influence the relative pathogen fitness within co-infection contexts. We found that an interaction between host sex and pathogen genotype determined relative pathogen fitness (Table 3). Upon examining each pathogen co-infection treatment individually, an interaction between sex and pathogen genotype influencing relative pathogen fitness was detected for every co-infection treatment except for sequential C24, C19 (Table 3). In females, the more virulent genotype (C19) produced the majority of transmission spores in each co-infection context except for when pathogen genotype C24 established first (Fig. 4). Conversely, co-infection in males resulted in equal spore production of the competing genotypes except for when the less virulent genotypes (C24 or C1) established before C19. In both of these cases, prior establishment of the less virulent genotype resulted in C19 being competitively inferior. In general, coinfections within females were represented by the disproportionate transmission of the more virulent pathogen, but this was not observed in males. For example, in sequential infections with C24, pathogen C19 produced 90.4% of the total spores within female hosts, but only produced 49.8% of spores within males.

**Figure 4:**
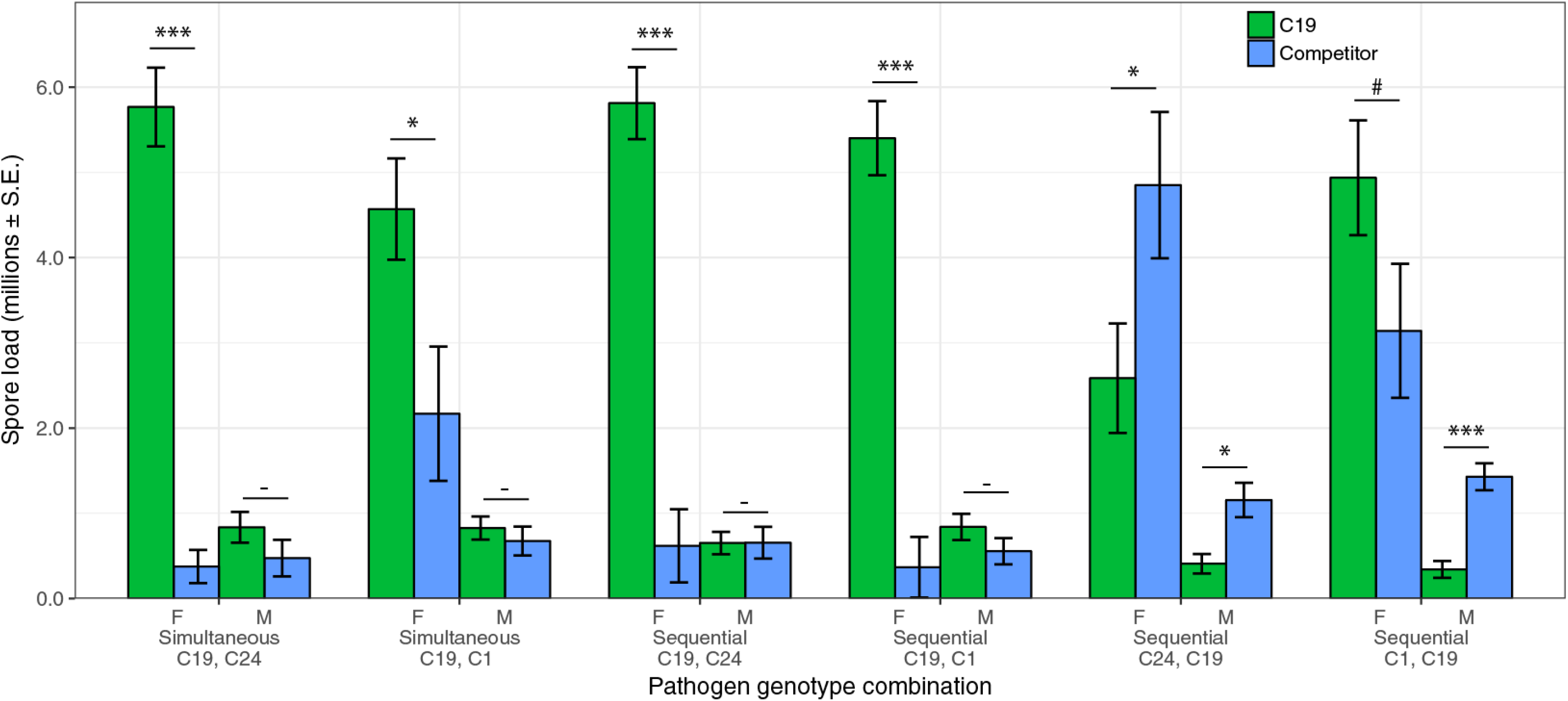
The influence of host sex and co-infection treatment on within-host pathogen competition as measured by relative spore production of individual genotypes. Green bars represent spore production by pathogen genotype C19 and blue bars represent spore production by the competing pathogen genotype (either C24 or C1). Shown are individual means for each genotype and standard errors. Asterisks indicate significant difference in mean spore production between competing genotypes within a single sex (two sample t-tests: *** *p* < 0.001, ** *p* < 0.01, * *p* < 0.05, # *p* < 0.10, – > 0.1

**Table 3:** Summary of an analyses of variance describing the effect of host sex, pathogen genotype, and their interaction on relative pathogen fitness within co-infection treatments. Asterisks denote significant effects (α = 0.05).

| Relative pathogen fitness<br>(relative spore production) | $\chi^2$ | d.f. | *p*-val |
| --- | --- | --- | --- |
| **Full model including all co-infection treatments** |  |  |  |
| Sex | 170.335 | 1,574 | <0.001 * |
| Pathogen genotype | 49.093 | 1,574 | <0.001 * |
| Sex : Pathogen genotype | 69.034 | 1,574 | <0.001 * |
| **Simultaneous C19, C24** |  |  |  |
| Sex | 40.511 | 1,72 | <0.001 * |
| Pathogen genotype | 57.442 | 1,72 | <0.001 * |
| Sex : Pathogen genotype | 43.890 | 1,72 | <0.001 * |
| **Sequential C19, C24** |  |  |  |
| Sex | 47.780 | 1,98 | <0.001 * |
| Pathogen genotype | 48.988 | 1,98 | <0.001 * |
| Sex : Pathogen genotype | 49.133 | 1,98 | <0.001 * |
| **Sequential C24, C19** |  |  |  |
| Sex | 26.541 | 1,88 | <0.001 * |
| Pathogen genotype | 6.991 | 1,88 | 0.008 * |
| Sex : Pathogen genotype | 1.776 | 1,88 | 0.183 |
| **Simultaneous C19, C1** |  |  |  |
| Sex | 29.257 | 1,84 | <0.001 * |
| Pathogen genotype | 6.960 | 1,84 | 0.008 * |
| Sex : Pathogen genotype | 5.386 | 1,84 | 0.020 * |
| **Sequential C19, C1** |  |  |  |
| Sex | 49.962 | 1,96 | <0.001 * |
| Pathogen genotype | 73.905 | 1,96 | <0.001 * |
| Sex : Pathogen genotype | 58.960 | 1,96 | <0.001 * |
| **Sequential C1, C19** |  |  |  |
| Sex | 42.688 | 1,106 | <0.001 * |
| Pathogen genotype | 0.538 | 1,106 | 0.463 |
| Sex : Pathogen genotype | 8.934 | 1,106 | 0.003 * |

## Discussion

Much of our understanding on how pathogens should exploit hosts and how exploitation strategies should evolve are influenced by studies focusing on single infections. Yet in nature, infection is more likely to co-occur between multiple pathogen genotypes of the same or different species (Read and Taylor 2001; Rigaud et al. 2010; Balmer and Tanner 2011). When multiple pathogens establish within a host, theory predicts that co-infection commonly favours more virulent pathogens through increased competition for host resources (Alizon et al. 2013). However, host heterogeneity may affect the relative fitness among competing pathogens (de Roode et al. 2004; Hodgson et al. 2004; Råberg et al. 2006; Ben-Ami et al. 2008; Ben-Ami and Routtu 2013; Izhar et al. 2015; Louhi et al. 2015) thus the evolution of more virulent pathogens is not necessarily a universal outcome of co-infection (Alizon et al. 2013; Cressler et al. 2016). In this study we considered how a common source of host heterogeneity in many species, the differences between the sexes in their capacity limit pathogen performance (see Table 1, Cousineau and Alizon 2014), can modify the expression of virulence in co-infections and the consequences this may have for the maintenance of genetic diversity in pathogen populations.

Our results indicate that the ability of co-infections to modify the relationship between pathogen growth (*i.e.* spore production) and pathogen induced reduction in host lifespan (*i.e.* virulence) will depend on the sex of the host. While infection in females is marked by a negative relationship between spore production and virulence, similar virulence patterns in male hosts correspond with more limited variation in spore production (Fig. 1). This was also observed in Thompson *et al.* (2017) where the scale of the trade-off between virulence and spore production varied due to host sex, driven largely by a reduction in overall variation in pathogen fitness across co-infection treatments in males. Despite this lower variation in spore production within male hosts, we found that the relative relationships between pathogen virulence and spore production were qualitatively similar between the sexes. Regardless of sex, simultaneous co-infections often exhibited relationships between pathogen virulence and spore production equal to that of the most virulent pathogen in isolation (Fig. 2, 3). Likewise, the relationship between virulence and spore production in sequential co-infections depended on pathogen genotype and order of establishment, but was unaffected by sex. These patterns culminate with females representing an environment within which pathogens can attain the highest levels of reproduction; yet coinfections within females also represent a substantial decrease in pathogen spore production as compared to the males.

Despite commonalities between the sexes in the overall patterns of transmission and virulence, the relative fitness between co-infecting pathogen genotypes varied strongly between males and females. We found that females exhibited significant differences in spore production between competing genotypes in all but one co-infection treatment, with the more virulent pathogen (C19) producing up to 5.3 million spores more than its competitor (Simultaneous C19, C24; Fig. 4). In contrast, when pathogens competed within males, each genotype produced an equal number of transmission spores in simultaneous exposures and sequential exposures when the more virulent pathogen established first (Fig. 4). Explaining these results may be that the more limited exploitative environment of male *Daphnia* (Thompson et al. 2017; Gipson and Hall 2018) prohibits pathogens from exhibiting variation in exploitation strategies. Males then may be a reservoir for pathogen genetic diversity, resulting in equal fitness between competing genotypes and even favouring genotypes which are frequently outcompeted within female hosts.

Our results also indicate that the arrival sequence of co-infecting pathogens will lead to different competitive outcomes depending on the host sex encountered. Previously established pathogens may inhibit later arriving competitors by blocking pathogen establishment, exhausting resources, or by inducing host immune responses (de Roode et al. 2005a; Lohr et al. 2010; Hoverman et al. 2013). In these situations, early establishing pathogens may exhibit considerable levels of reproduction even in competition with more virulent genotypes. Indeed, Ben-Ami *et al.* (2008) found that less virulent genotypes can exhibit substantial levels of reproduction when establishing first even though they are outcompeted when establishing after more virulent genotypes. Here, females exhibited this general pattern with the more virulent C19 genotype outcompeted by C24 and producing a similar number of spores as C1 when establishing second (Fig. 4). Conversely, the constraints imposed by male hosts appear to keep pathogen C19 from ever outcompeting other genotypes when they establish first. Consequentially, male hosts may thus maintain pathogen genetic variation that would otherwise be eroded by infection in females where the more virulent genotype is more frequently transmitted.

Taken together, our results suggest that the evolutionary outcome of pathogen virulence in coinfection contexts will depend on how often pathogens encounter male and female hosts. In coinfections, more virulent pathogens often transmit more than their less virulent competitors, potentially favouring the evolution of higher virulence (de Roode et al. 2005b; Bell et al. 2006; Råberg et al. 2006; Ben-Ami et al. 2008; Balmer et al. 2009; Ben-Ami and Routtu 2013). Here, we find that co-infections in female hosts favour the transmission of the more virulent pathogen, potentially selecting for higher levels of virulence. In contrast, co-infection in males is characterized by either equal fitness of competing pathogens or higher fitness of less virulent pathogen genotypes. Co-infection in males may then either slow the evolution of virulence or completely reverse selection on virulence depending on the exposure context. Yet, the evolutionary outcome of virulence will depend not only on the within-host outcomes of infection, but also in how disease is transmitted between hosts (van Baalen and Sabelis 1995; Frank 1996; Mideo et al. 2008; Choisy and de Roode 2010; Alizon et al. 2013). Indeed, sex ratio can vary within a variety of species (Clutton-Brock and Iason 1986; Duneau and Ebert 2012 and Table 2 therein), including *Daphnia* (Galimov et al. 2011), which may influence the frequency of pathogen transmission between each sex and thus the overall patterns of selection on virulence. A higher proportion of the more exploitable sex, for example, may favour the transmission of more virulent pathogens, whereas population shifts to the less exploitable sex may restrain or reverse this pattern.

The ubiquity of pathogens in natural populations suggests that individuals are likely to encounter and become infected by multiple pathogen genotypes. We show that host sex may further impact on the relative fitness among pathogen genotypes due to within-host competition, finding that differences in the scale of the trade-off between pathogen virulence and transmission between the sexes affect the relative fitness among competing pathogens. This work reaffirms the role that host heterogeneity can play in affecting the outcome of within-host pathogen competition and its potential to influence the evolution of virulence (de Roode et al. 2004; Hodgson et al. 2004; Råberg et al. 2006; Ben-Ami et al. 2008; Ben-Ami and Routtu 2013; Izhar et al. 2015). Ultimately, how virulence should evolve within co-infection contexts will depend on frequency of encountering each sex as well as variation in the exploitative potential of male and female hosts. Collectively, our results suggest that the often observed relationship between virulence and pathogen competitive ability may not apply equally to each sex, providing a mechanism for the maintenance of pathogen genetic variation in sexually dimorphic host populations.

## Acknowledgements

We would like to thank Isobel Booksmythe, Toby Hector, Louise Nørgaard, and Kyle Kelly for helpful conversations, laboratory assistance, and advice on statistical analysis.

